# Synthetic observations from deep generative models and binary omics data with limited sample size

**DOI:** 10.1101/2020.06.11.147058

**Authors:** Jens Nußberger, Frederic Boesel, Stefan Lenz, Harald Binder, Moritz Hess

## Abstract

Deep generative models can be trained to represent the joint distribution of data, such as measurements of single nucleotide polymorphisms (SNPs) from several individuals. Subsequently, synthetic observations are obtained by drawing from this distribution. This has been shown to be useful for several tasks, such as removal of noise, imputation, for better understanding underlying patterns, or even exchanging data under privacy constraints. Yet, it is still unclear how well these approaches work with limited sample size. We investigate such settings specifically for binary data, e.g., as relevant when considering SNP measurements, and evaluate three frequently employed generative modeling approaches, variational autoencoders (VAEs), deep Boltzmann machines (DBMs) and generative adversarial networks (GANs). This includes conditional approaches, such as when considering gene expression conditional on SNPs. Recovery of pair-wise odds ratios is considered as a primary performance criterion. For simulated as well as real SNP data, we observe that DBMs generally can recover structure for up to 100 variables with as little as 500 observations, with a tendency of over-estimating odds ratios when not carefully tuned. VAEs generally get the direction and relative strength of pairwise relations right, yet with considerable under-estimation of odds ratios. GANs provide stable results only with larger sample sizes and strong pair-wise relations in the data. Taken together, DBMs and VAEs (in contrast to GANs) appear to be well suited for binary omics data, even at rather small sample sizes. This opens the way for many potential applications where synthetic observations from omics data might be useful.

## General Background

Tabular categorical data occur frequently in biomedical research. Examples are single nucleotide polymorphism (SNP) data, where a large number of SNPs is measured for each of a number of individuals. While such measurements typically are linked to a phenotype or an outcome, such as survival, the structure of the data itself, such as the co-occurrence of SNPs also is of interest. Deep generative models build on neural network architectures and can learn the corresponding joint distribution of such data. Popular examples are deep Boltzmann machines (DBMs, [1]), variational autoencoders (VAEs, [2]) and generative adversarial networks (GANs, [3]).

There are many successful biomedical applications based on learning the joint distribution of data. For example in [4], the authors demonstrated how GANs can be employed to simulate biomedical data to facilitate data analysis under privacy restrictions. Using GANs, Yelmen et al. [5] generated artificial chromosomes from real SNP data. Lopez et al. [6] employed VAEs for learning a low dimensional representation of gene expression in single cells and in [7], the authors performed drug response prediction with VAEs. DBMs have been successfully applied in learning the joint distribution of SNP data [8].

After training, the above described models can generate synthetic observations with similar properties as observed in the data used for training. This property has, e.g., been used for denoising single cell gene expression data [9]. Another major challenge that can be addressed by deep generative models is data privacy [10]. Specifically, deep generative methods have been proposed to generate synthetic observations from the learnt joint distribution of the data [11], [4]. These synthetic observations might then be securely shared.

A potential major hurdle in all these promising use cases for deep generative models is limited sample size, as deep learning techniques frequently have been developed with large sample sizes in mind. It is still unclear whether reasonable performance can be expected when the number of measured features is relatively large compared to the number of observations. Therefore we systematically evaluate the performance of the above mentioned deep generative approaches, DBMs, VAEs and GANs, with a specific focus on binary data, e.g., as relevant for SNP data. We consider simulated data as well as SNP data from the 1000 genomes project [12], to compare the approaches. The results of this investigation should then provide guidance for selecting the appropriate approach for a specific available sample size.

While it is generally considered difficult to judge the quality of synthetic data with a high number of dimensions in terms of their distance to the training data [13], we employ simple summary statistics, specifically odds ratios (ORs) to investigate the distribution of pair-wise correlations. Compared to a measure for overall similarity, like the Euclidean distance between two data-points, this measure is more robust and easier to interpret while giving an estimate for the lower-bound of the similarity of the joint-distributions of generated, synthetic data and the training data.

Since deep generative approaches also can model the conditional distributions of different data modalities, e.g. for gene expression conditional on SNP data [14], we also evaluate such conditional approaches, specifically conditional GANs (cGANs; [15]), and compare these with DBMs which natively allow to model conditional distributions.

In the following, we provide a brief overview of deep generative approaches, before evaluating their performance first with a single data-type (simulated and real SNP data) and then with simulated data from a conditional scenario. We close with general recommendations for the use of deep generative approaches with rather small sample size. To facilitate subsequent use of the approaches in such settings, a Jupyter notebook with further examples is provided as a supplement.

## Deep generative models

All investigated deep generative models do more or less build on the concept of artificial neural networks (ANN). The name is derived from the analogy to neurons in living organisms. Neurons, hereafter called units, transduce a signal to other cells when a certain signal threshold is reached. The strength of the signal in one cell is modeled as a linear combination of the signals from all connected units. In ANNs this linear combination is usually transformed using a non-linear activation function such as the sigmoid function, keeping differentiability for means of numerical optimization. Having many of these linear combinations with subsequent non-linear transformations in parallel, allows to model complex, non-linear dependencies in the data. VAEs and GANs incorporate feed-forward neural network where units are arranged in fully connected layers of hidden units with directed connections. The modeled networks thus have an input layer and an output layer with a directed signal path without loops. Having an unambiguous mapping from input to output, the feed-forward neural network is modeled as a parameterized function. The parameters correspond to the connection weights between layers.

Let *θ* = (*W*^(1)^, *W*^(2)^, *b*^(1)^, *b*^(2)^) be the set of parameters describing the interconnection of units for a feed-forward neural network with one hidden layer. Matrices *W* contain the weights of the interconnections between layers and vectors the bias terms *b*. Given a vector *x* as input and a non-linear activation function *f*, the output *y* is then calculated as

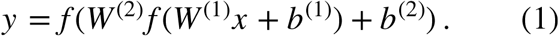

### Variational autoencoder (VAEs)

Autoencoders are feed-forward neural networks that have a reconstruction 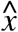 of the input *x* as output. The parameters are optimized with respect to this reconstruction. A rather small number of hidden units in an intermediate layer acts as a bottleneck that enforces reduction to the most important properties of the data. This approach therefore learns a lower dimensional representation (encoding) of the data, from which the model can reconstruct (decode) the original data. If the distribution of encodings is known and the autoencoder generalizes the original data distribution well, one could explore the modeled distribution by drawing from the encodings. However, the true underlying distribution is unknown and cannot be inferred from the training data. However, if we attempt to approximate the latent distribution of encodings by variational inference techniques, we arrive at a variational autoencoder (VAE) [2] (see Figure 1, Panel B). Here the low dimensional intermediate layer comprises stochastic, units (*z*).

**Figure 1:**
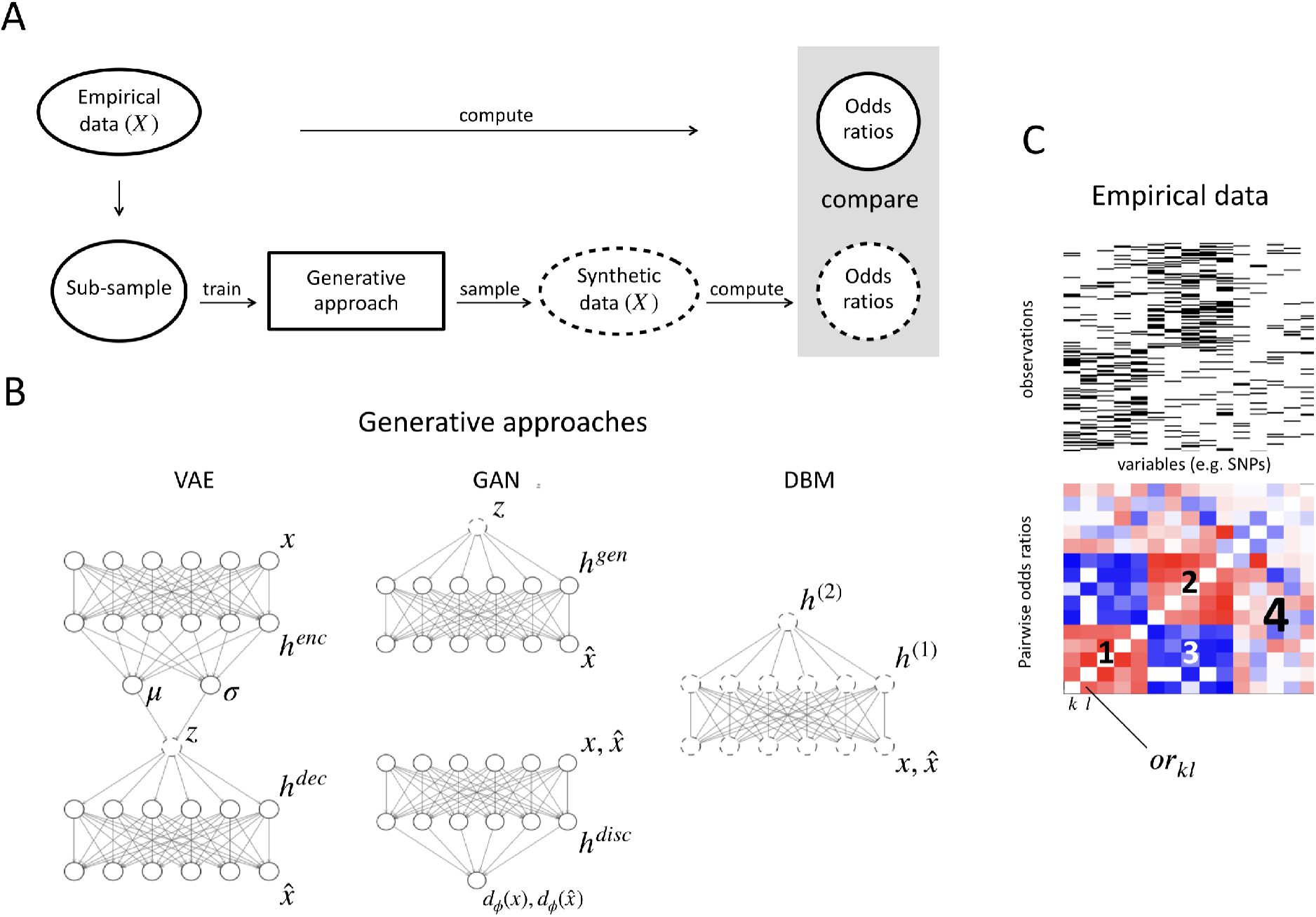
(A) Experimental design used to compare VAEs, DBMs and GANs. A data-set of tabular binary data (empirical data) is sub-sampled and used to train a deep generative model. The performance of the three deep generative approaches is then judged by comparing pairwise odds ratios computed from the empirical data and the synthetic data, drawn from the trained models. (B) Schematic architecture of the deep generative approaches (h = hidden layer, enc = encoder, dec = decoder, gen = generator, disc = discriminator). Solid circles indicate deterministic units and dashed circles indicate stochastic units. (C) Exemplary data and odds ratios. There are three different groups of variables. Odds ratios within and between groups are considered, resulting in four blocks of odds ratios. Odds ratios are shown log transformed. Red indicates positive log odds ratios while blue indicates negative log odds ratios.

Let *q_ϕ_* denote the approximation of the conditional distribution of the latent representation given the input, implemented via the encoding neural network and *p_θ_* the distribution corresponding to the observed data, e.g. implemented for the conditional distribution given the latent representation by the decoding neural network. During optimization, we maximize the likelihood of *N* individual data-points, given as

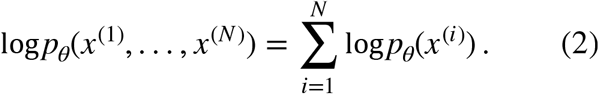

Since we specify an approximate (prior) distribution for the latent representation in a variational inference framework, we can only maximize the likelihood up to the lower bound 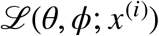, given as

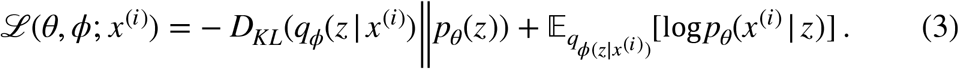

During training, the first right-hand term penalizes the deviation of the distribution of the latent encodings from the prior distribution, as indicated by the Kullback-Leibler divergence (*D_KL_*), while the second terms indicates, how well the input can be reconstructed, given the latent encodings.

With the assumptions of having normally-distributed encodings *z* and a Bernoulli-distributed output layer, the lower bound, shown in Equation 3, is given by

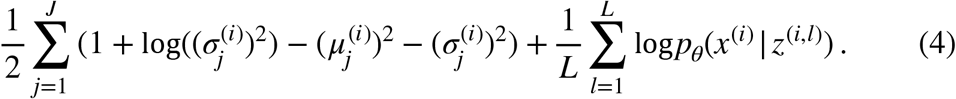

*μ* and *σ* relate to the mean and standard deviation of the normal distribution, from which the encodings are drawn. *μ*^(*i*)^ and *σ*^(*i*)^ result from propagating the input *x*^(*i*)^ through the neural network to the units *μ* and *σ* (see Figure 1, Panel B). *z*^(*i,l*)^ is drawn according to 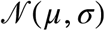. *J* denotes the dimensionality of the latent encodings and *L* is the number of encodings, drawn to approximate the expectation.

Drawing from the modeled distribution, i.e. generating synthetic observations, now is straightforward by drawing from the normal distribution prior and propagating the result through the decoding network. The role of the decoder will later also be referred to as a “generative function”.

### Generative Adversarial Networks (GANs)

As described for the VAEs, a generative function can be used to transform realizations *z* coming from some source distribution to realizations 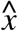 of a distribution that matches the distribution of the training data *x*. While this source distribution is motivated by variational inference in VAEs, generative adversarial networks (GANs) use an informative source distribution. In addition to the generating function, which should transform samples from the source distribution into samples resembling the original data at hand, a second feedforward neural network, the discriminator, is trained as an adversary that discriminates between original data (*x*) and samples stemming from the generator 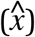 (see Figure 1, Panel B). The corresponding loss function can be used to optimize both the discriminator and the generator. As the performance of the discriminator increases, the generator is optimized so that it will continuously fool the discriminator. This back and forth is implemented in the training process as a min-max game, optimizing one network while the other serves the gradient that is used for optimization.

Let *g_ϕ_* denote the generator, *d_ϕ_* the discriminator, *P*_data_(*x*) the empirical distribution over the data and *p_z_* the source distribution from which *z* is drawn. The min-max game between *g* and *d* can then be stated as

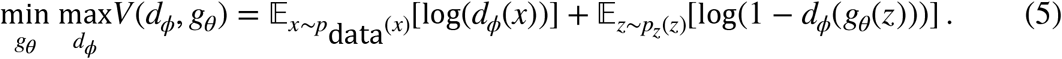

After training, synthetic observations can be generated by initializing *z* to a random state and propagating the information through the generator network. In the binary example, the resulting real valued activations are then transformed to binary values 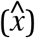 by employing a hard threshold at 0.5.

GANs can also be used in a conditional setting, where the model generates data based on a random vector and a given condition *c*, resulting in cGANs. For example, *c* could incorporate SNP information when synthetic gene expression data should be generated conditional on SNP status. The additional conditional information is provided to the generator and the discriminator during training.

During sampling, *c* is presented together with a random vector *z*, yielding generated data, that mimics the distribution of the training data respecting the condition *c* (for further applications, e.g. see [16] and[17]).

### Deep Boltzmann Machines (DBMs)

While VAEs and GANs incorporate feed-forward networks, which are used to transform random variables, into realistic synthetic observations, and intermediate hidden layers only specify the transformation, in deep Boltzmann machines, all hidden units (as well as the observation level, called visible units) are considered to be random variables. In our context of binary data, the units can take on two states, zero and one. Connections between units are not directed and the input (*x*) layer at the same time also is the output layer 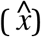 (see Figure 1, Panel B). Latent variables, arranged in hidden layers (*h*), can capture non-linear dependencies between the observed variables, similar to VAEs and GANs.

Deep Boltzmann machines are derived from Boltzmann machines [18] which are closely related to an energy model of particles explored in physics, where a likely configuration of particles should result in a low energy and vice versa. For a DBM with two hidden layers and parameters *θ* = (*W*^(1)^, *W*^(2)^, *b*^(0)^, *b*^(1)^, *b*^(2)^), we have the energy function

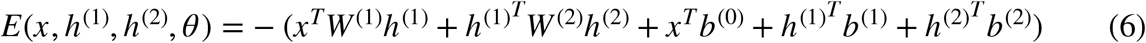

which indicates the energy for a configuration of the states of hidden and observed nodes. By normalizing through the partition function *Z*, the energy is transformed into a probability as

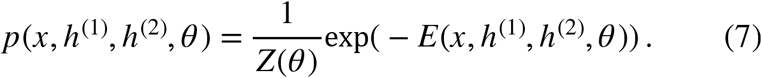

*Z* here is the sum over all possible configurations of the states in observed and hidden units:

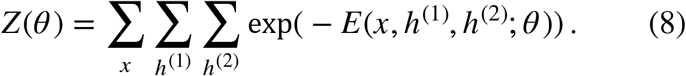

In the training procedure the likelihood of the training data *x*, given as

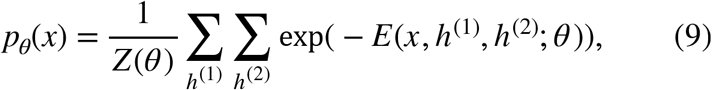

is maximized. After training, given a random initialization of the model parameters, we can run a Markov chain between *x* and in this example *h*^(1)^ and *h*^(2)^. When we arrive at an equilibrium after a finite number of steps, where the Markov chain converges, we retrieve a synthetic observation 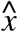 from the model. This procedure is also called Gibbs-sampling.

As it is difficult to jointly optimize the states of all latent and observed variables in a DBM with multiple hidden layers, parameters of DBMs typically are optimized in a two-step technique [1]. In a first step, shallow DBMs, comprising only two layers, also called restricted Boltzmann machines (RBMs, [19]) are consecutively trained. In a second step, the parameters of a stack of RBMs are jointly optimized.

DBMs also can provide information on conditional distributions. During training, the conditional input is provided in distinct additional units in the visible layer, i.e. *x* = (*x_nc_, x_c_*), with *x_nc_* being the variables whose conditional distribution is to be learned and *x_c_* being the condition. The DBM then will be trained to learn the joint distribution of *x*. When sampling from the trained DBM, samples from the conditional distribution *p*(*x_nc_* | *x_c_*) are obtained by fixing the visible units that correspond to the condition during the Gibbs-sampling procedure, meaning that after each step *t* of the Markov chain, in the given sample, *x*^(*t*)^, consisting of (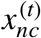 and 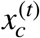), 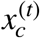 is set back to the condition *x_c_*.

## Material and Methods

### Simulated SNP Data

We first evaluate the performance of the three deep generative approaches based on binary simulated data which roughly reflect the structure of SNP haplotypes. A haplotype consists of 50 SNPs and the two states correspond to the major and minor alleles respectively. In the conditional simulation setting, we investigate 10 SNPs which can affect the (dichotomized) expression of 10 out of 50 genes, totaling to 60 variables.

In the non-conditional setting, there are two types of individuals occurring by equal chance of 0.5. In the first type, the first five features take on the value 1 with a probability of 0.37 and the features six to fifty with a probability of 0.1. In the second type, the variables 6 to 10 take on the value 1 with a probability of 0.37 while this probability is 0.1 for the remaining variables. In our interpretation, the chance of 0.37 represents a signal, while 0.1 is a uniformly distributed noise. An example for the structure of the data is shown in Panel C of Figure 1. When computing pair-wise odds ratios from the above described data, four blocks of odds ratios emerge. Within block 1 and 2, there is an on average positive log odds ratio of 0.53, in block 3 there is an on average negative log odds ratio of –0.62 and in block 4 the log odds ratio is around 0, corresponding to the absence of any structure (see Figure 1, Panel C). Given a data-set of 10.000 simulated haplotypes comprising 50 SNPs, sub-samples of 100, 1000 and 5000 observations are drawn and employed to train a generative model. Afterwards 200 samples are drawn from trained models to compute log odds ratios (see Figure 1, Panel A). The whole procedure is repeated 200 times and averaged results are considered.

In the conditional setting, there also are two types of individuals for the SNP data, occurring with equal chance of 0.5. In a non-information carrying mode, a SNP variable is simulated with a chance of 0.15 to take on the value 1. The corresponding gene expression data is independently simulated with variables having a chance of 0.15 to take on the value 1. This type can be interpreted as random non-interacting noise between the SNP and gene expression data and results in no association between the variables, corresponding to a log odds ratio of 0. The second type of individuals for the SNP data, denoted as information inducing mode, induces an association between the 10 SNPs and the genes one to ten. *SNP*_1_ is capable to induce high or low expression in *Gene*_1_, *SNP*_2_ in *Gene*_2_ and so on. The information inducing mode again has two types of individuals occurring with equal chance of 0.5. In the first type, each SNP takes on the value of 1 with a probability of 0.05, in the second type, a SNP takes on the value of 1 with a probability of 0.95. Both times, the probability of a gene to be highly expressed, i.e. having a value of 1 is *p*(*gene*_1_ = 1| *SNP_i_* = 1) = 0.63. This means that in the information inducing type, a dependency between the SNP data and the gene expression data is induced, in contrast to the non-information inducing type. On average, the log odds ratio between SNPs and gene expression is 1.1. An example for the conditional data is shown in Supplementary Figure 1.

### 1000 genomes SNP data

Single nucleotide polymorphisms (SNPs) in the HLA-B gene locus were defined by the NCBI 1000 genomes browser, using the GRCh37.p13 genome version. Phased haplotypes were retrieved from phased 1000 genomes data as provided for the IMPUTE software (https://mathgen.stats.ox.ac.uk/impute/1000GP_Phase3.html). In total 5008 haplotypes are analyzed, belonging to 2504 individuals. All variants with a minor allele frequency (MAF) below 0.05 and above 0.2 were discarded, resulting in 110 SNPs that are investigated.

### Training of deep generative models

We intend to compare the three deep generative modeling approaches “out of the box”, meaning in a default configuration. Therefore we employ simple parameter settings for the three deep architectures using only one to two latent layers. As we are modeling Bernoulli distributed, binary data, we use the sigmoid function as activation function for the units modeling the observed variables. For the latent variables, we employ sigmoid activation functions in the DBM, the Tangens hyperbolicus (tanh) in the VAE and the leaky rectified linear unit (ReLu) in the GAN. VAEs, GANs and cGANs can be trained by backpropagation of errors, so there is a multitude of frameworks available such as TensorFlow [20]. We employ the Flux package [21] written for the Julia programming language [22] to fit these approaches using the parameter settings as described in the supplementary information.

In contrast, the training procedure of DBMs differs fundamentally from the latter. Here Markov chain Monte Carlo (MCMC) techniques are employed to estimate the states of latent and observed variables and these techniques cannot easily be implemented using the aforementioned frameworks. In addition, the training procedure of DBMs involves two steps, a layer-wise pre-training and a fine-tuning step. In the pre-training step, the weights between two adjacent layers are initialized to meaningful values and in the fine-tuning step, all weights are jointly fine-tuned. In general we follow the procedure as described in [8]. In order to compare the DBM with the GAN and VAE, we perform layer-wise pre-training until we observe a minimum for the reconstruction error. The results for different training status during the joint refinement of the parameters are then compared with the results from GAN and VAE. Hyper-parameter settings are shown in the supplement. We fit DBMs with the BoltzmannMachines package [23] provided for the Julia programming language.

### Sampling from deep generative approaches

We generate synthetic observations by initializing the stochastic variables in the models to random states and then propagating the information through the decoder network (VAE), or the generator network (GAN). In the DBM we run a Gibbs chain for 100 steps. Since in the VAE and the GAN the output layer 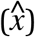 consists of deterministic units, a hard threshold is applied at 0.5 to transform the real valued output of the employed sigmoid activation function to a binary output.

### Evaluation of deep generative approaches

Evaluation of deep generative models is an active field of research and still lacks broadly applicable solutions [13]. While all models might be investigated based on the loss which is optimized during training, this kind of measure is not useful for comparing different generative approaches.

As an alternative, we propose to employ summary statistics raised over the generated synthetic observations. In particular in medical research, where privacy issues impose regulations on data processing and publication, the use of summary statistics has proven to be effective [24].

Specifically, we propose to use odds ratios (see [25]), calculated between the features in the data. Odds ratios indicate the degree of co-occurrence of binary features based on frequencies in a cross table. If some of the cells in the cross table happen to be zero, they are replaced by the value 0.5 for computational reasons.

Calculating odds ratios between all *p* features in the data results in a symmetric matrix 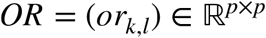. For convenience of calculation and interpretation, all computed odds ratios are transformed by the logarithm (log) function. The advantage of logarithmic odds ratios is that their distribution is roughly symmetric [25] which simplifies further analyses.

For the experiments with simulated data, the *OR* matrix is computed from 200 data-sets, each comprising 200 synthetic observations sampled from the generative models. For each data-set the logarithmic pair-wise odds ratios are computed, resulting in the matrices (*OR*^(1)^,…, *OR*^(200)^). The final statistic, shown in the figures is the mean over each of the entries in the *OR*^(*k*)^ matrices, resulting in a matrix 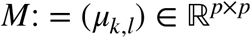 of means with 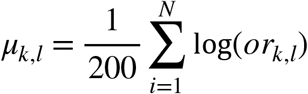.

In the conditional setting, where we aim at generating the (binary) expression status of genes based on SNP data, *x_l_* represents a SNP and *x_k_* represents a gene.

In the application with real SNP data, the focus is less on quantifying the performance for a single variable, i.e. a SNP, but we are rather interested in the overall performance. Consequently, the performance of the generative models is judged by a measure which quantifies the overall deviation from the empirical data. Specifically the mean squared error (MSE) of the difference between all pair-wise log odds ratios computed from the synthetic data (samples) and test data, not used for training, is employed.

## Results

### Non-conditional setting - simulated data

In the following we investigate, how strong log-odds ratios in the blocks 1, 3 and 4 of the simulation design deviate from the log odds ratios in the training data and in the total data-set that was sub-sampled (see Figure 1).

At a training data size of 100 observations, we observe that there is already a large amount of error in the sub-sample used for training, relative to the full data-set (Figure 2 and Supplementary Figure 2, upper row, green dots vs. green line). Specifically, log odds ratios computed from the training data, are on average far less extreme compared to the true log odds ratios in the full data-set. At this sample size, the performance of the three approaches differs considerably. While the log odds ratios learned by the VAE matches the log odds ratios in the training data well, there is a large spread in the log odds ratios learned by the DBM. However, when investigating the log odds ratios in block 4, i.e. between variables without any dependency, we observe that the VAE does generally under-estimate the log odds ratios, while the DBM accurately learns the distribution of log odds ratios in this block (Supplementary Figure 3). In contrast, the GAN is not able to pick up a signal, especially in the information carrying blocks 1 and 3.

**Figure 2:**
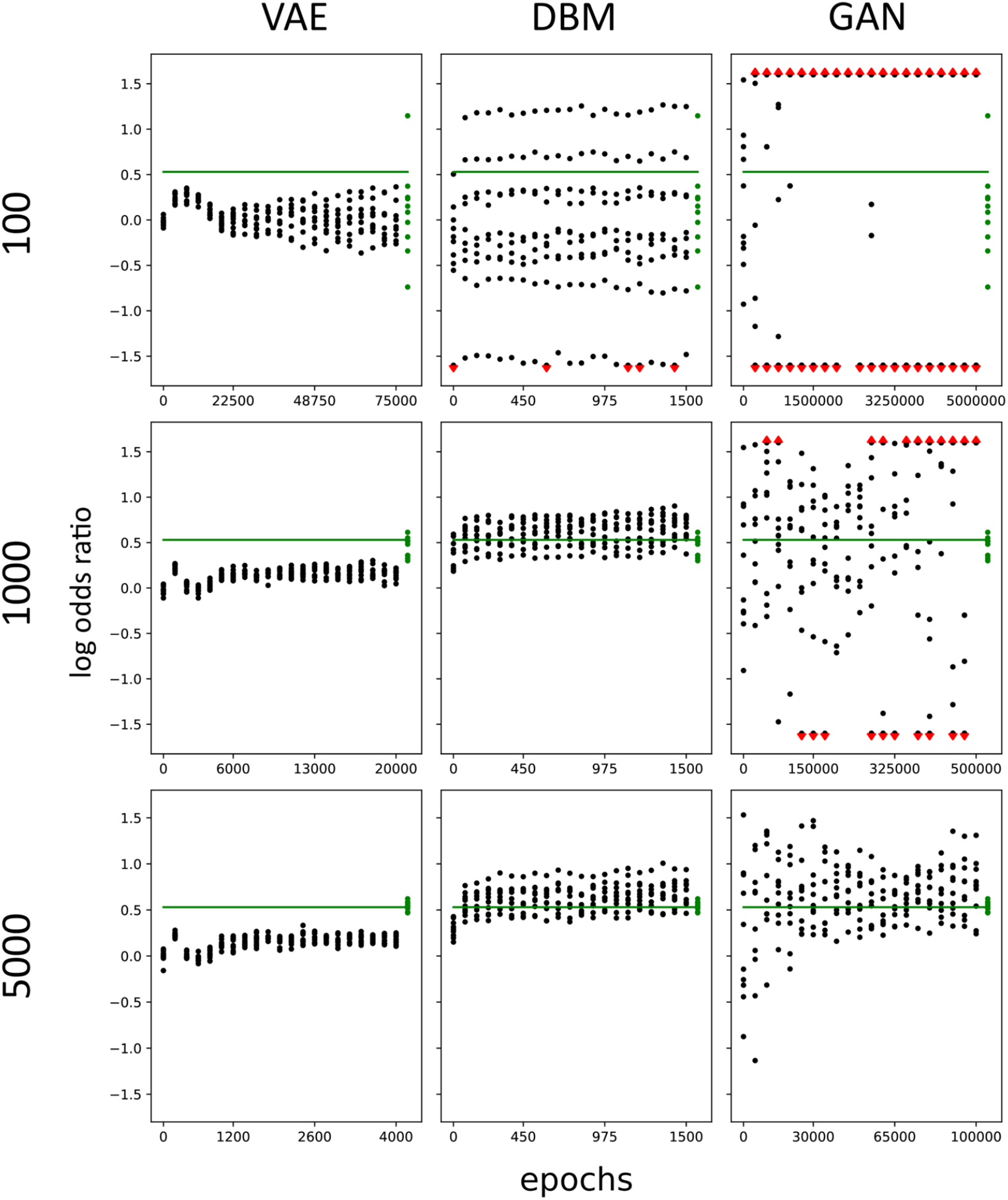
*Log odds ratios learned by the VAE, DBM and GAN approaches from simulated data. VAEs, DBMs and GANs are trained on* 100, 1000 *and* 5000 *observations of simulated SNP haplotypes in 50 SNPs. Log odds ratios between positively correlated SNPs (block 1, see Figure 1) are calculated* 20 *times during training. Each dot represents the average of the log odds ratios between two SNPs computed from synthetic observations, sampled from 200 generative models that were trained on different random sub-sets. The green line indicates the value of the theoretical truth according to the data generation procedure and the green dots indicate the actual values for the log odds ratios computed from the training set. Red arrows indicate values that are higher / lower than a log odds ratio of 1.5/-1.5*.

With increasing training data size, the log odds ratios computed from the training data match the true log odds ratio better (Figure 2, middle and lower row). At a training data size of 1000 observations, the VAE displays more stable results over the inspected epochs during training, while the strength of the log odds ratios are still under-estimated in the non-information carrying block 4 (Supplementary Figure 3). In contrast to the small training data set of 100 observations, this under-estimation is now also observable for the information carrying blocks 1 and 3 (Figure 2, Supplementary Figure 2). We do not observe a clear reduction of the under-estimation of the average log odds ratios between using 1000 and 5000 observations for training. In contrast to the VAE, the DBM rather over-estimates the log odds ratios in the information carrying blocks 1 and 3 (Figure 2 and Supplementary Figure 2). Here, we also do not observe a clear improvement from 1000 to 5000 observations. The GAN does only learn the log odds ratios in the information carrying block 1 with a performance that is comparable with the DBM at a training data size of 5000 observations (Figure 2). In block 3 the performance is worse (Supplementary Figure 2). In addition, the range of the log odds ratios is largely over-estimated in the non-information carrying block 4 (Supplementary Figure 3).

In order to investigate how much the results differ, when employing different training data, we also investigate the performance using three additional sets of 10.000 simulated observations. At 5000 sub-sampled observations, used for training, we observe stable results for the VAE and DBM over the investigated four data-sets, while the performance of the GAN is more dependent on the particular data-set (Supplementary Figure 4).

The behavior described above is also visually observable in the synthetic observations, generated from the models. At all sample sizes, the simulated pattern in the training data (Figure 1, Panel C) is not visually apparent in the observations drawn from the VAE. In contrast, the DBM and the GAN (at 5000 training examples) are able to generate data that resembles the empirical training data (Figure 3).

**Figure 3:**
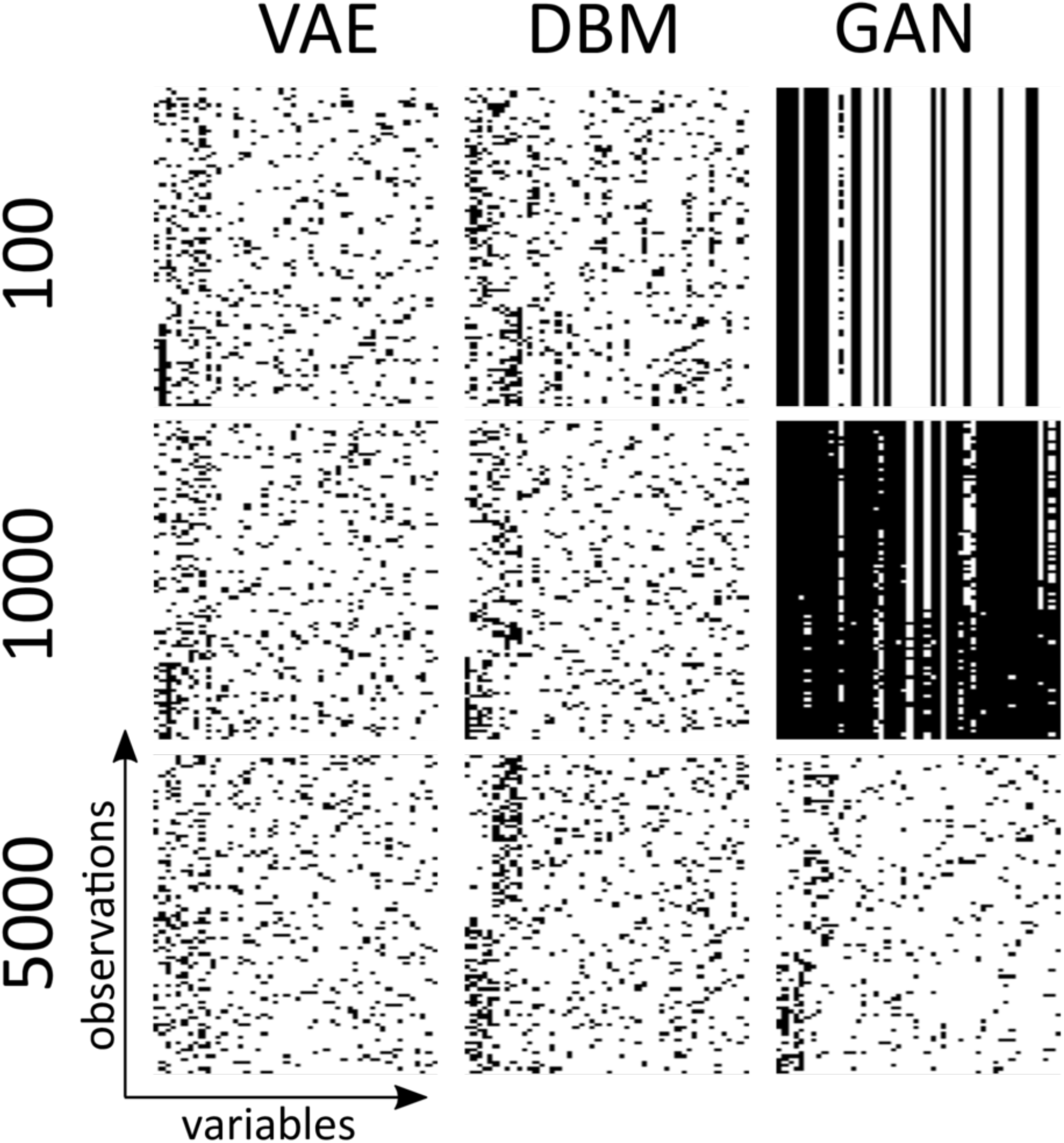
*Exemplary synthetic observations sampled from deep generative approaches. Random examples of 100 synthetic observations generated by trained VAEs, DBMs and GANs are shown. The generative models were trained on sub-sets comprising* 100, 1000 *and* 5000 *simulated observations. We always used the last state in the training process, meaning the samples correspond to models that were investigated at step 20 during the training process (see Figure 2). Synthetic observations are clustered based on their overall similarity. Black indicates a 1 while white indicates a 0*.

### Non-conditional setting - SNP data from HLA-B locus

In order to evaluate how well the investigated methods learn the joint distribution of SNPs in real data, we investigate haplotypes in the HLA-B gene locus which is involved in immune responses (https://ghr.nlm.nih.gov/gene/HLA-B). We use the 1000 genomes data [12], comprising 5008 haplotypes and inspect haplotypes of 110 SNPs (see Supplementary Figure 5 for haplotypes and pair-wise log odds ratios). As for the simulated data, we test, how well the approaches capture the pair-wise log odds ratios, i.e. the linkage disequillibrium (LD) pattern, found in the HLA-B locus. To investigate the performance, conditional on the size of the training data, we investigate the performance on subsets of 500, 1000 or 2000 randomly selected haplotypes. Since we aim at mimicking a real biomedical application, we do not investigate the learning of log odds ratios over multiple epochs but instead tune the number of training epochs conditional on the random initialization of the network parameters. To that end we train the models for several epochs (exact numbers are provided in the supplementary material), draw 1000 samples, i.e. synthetic data from the trained models, 20 times during training and compute the pairwise log odds ratios for the drawn samples and test data, not used for training (36.8% of the 5008 haplotypes). We then investigate the mean squared error (MSE), i.e. the aggregated difference between the log odds ratios in the test data and the sampled, synthetic data, and here report the results from the epoch with lowest MSE.

Compared to the VAE and DBM, the GAN performs worse in capturing the rather complex structure in the HLA-B locus. This is reflected in a comparably worse MSE (Figure 4), observable on the level of individual log odds ratios (Figure 5) and sampled haplotypes (Supplementary Figure 6), especially at the lower sample sizes of 500 and 1000 haplotypes. Even at 2000 haplotypes used to train the GAN, a large amount of log odds ratios are not well learned.

**Figure 4:**
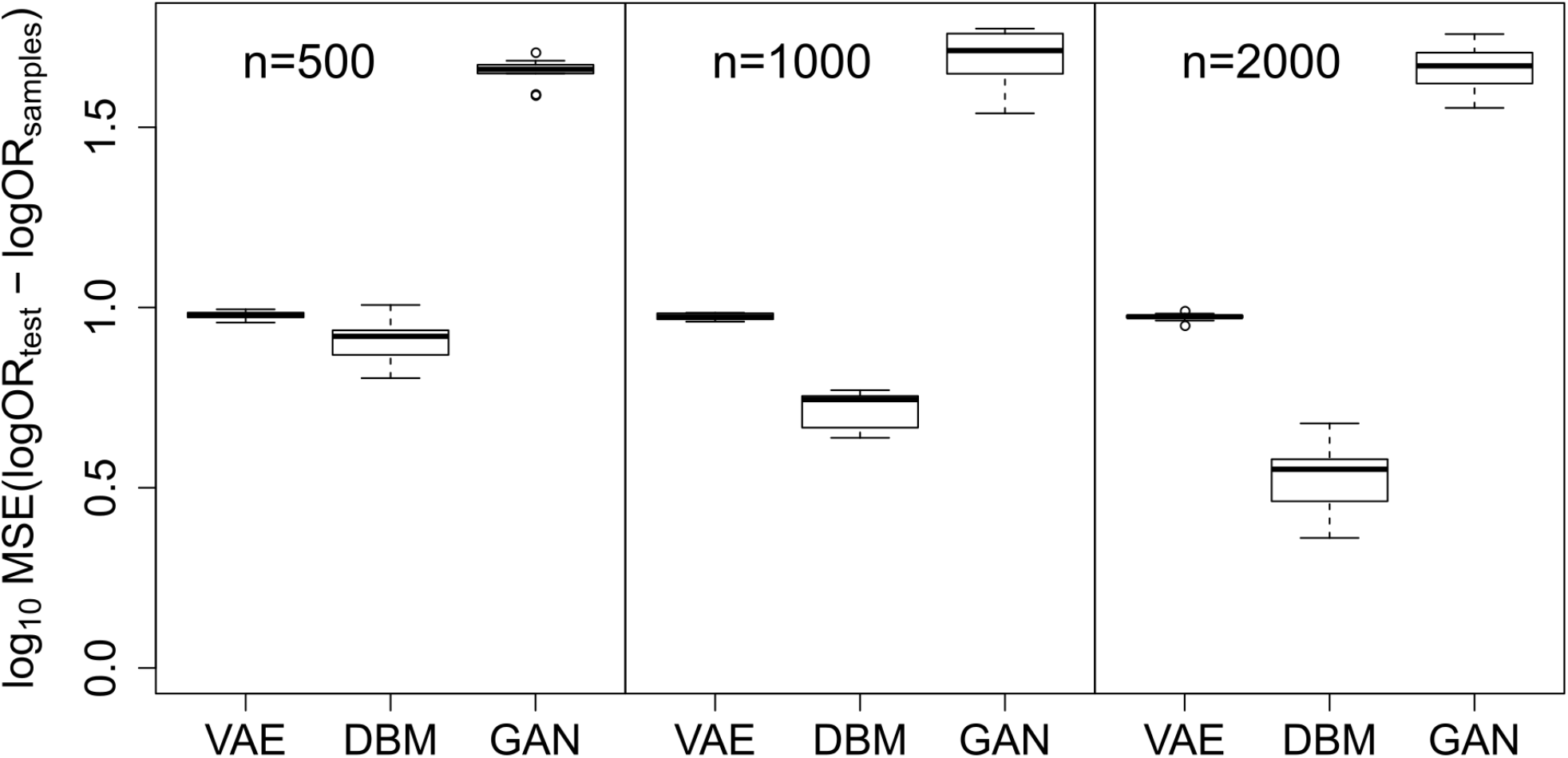
*Differences between log odds ratios computed from synthetic data and log odds ratios computed from real HLA-B data. Figure shows the mean squared error (MSE) between pair-wise log odds ratios computed from synthetic observations, sampled from VAEs, DBMs and GANs trained with* 500, 1000 *and* 2000 *haplotypes (samples) and log odds ratios computed from 36.8% of the 5008 haplotypes, not used for training (test). The shown variation is due to 10 randomly drawn sub-sets from in total 5008 available haplotypes and different random initializations of the model parameters*.

**Figure 5:**
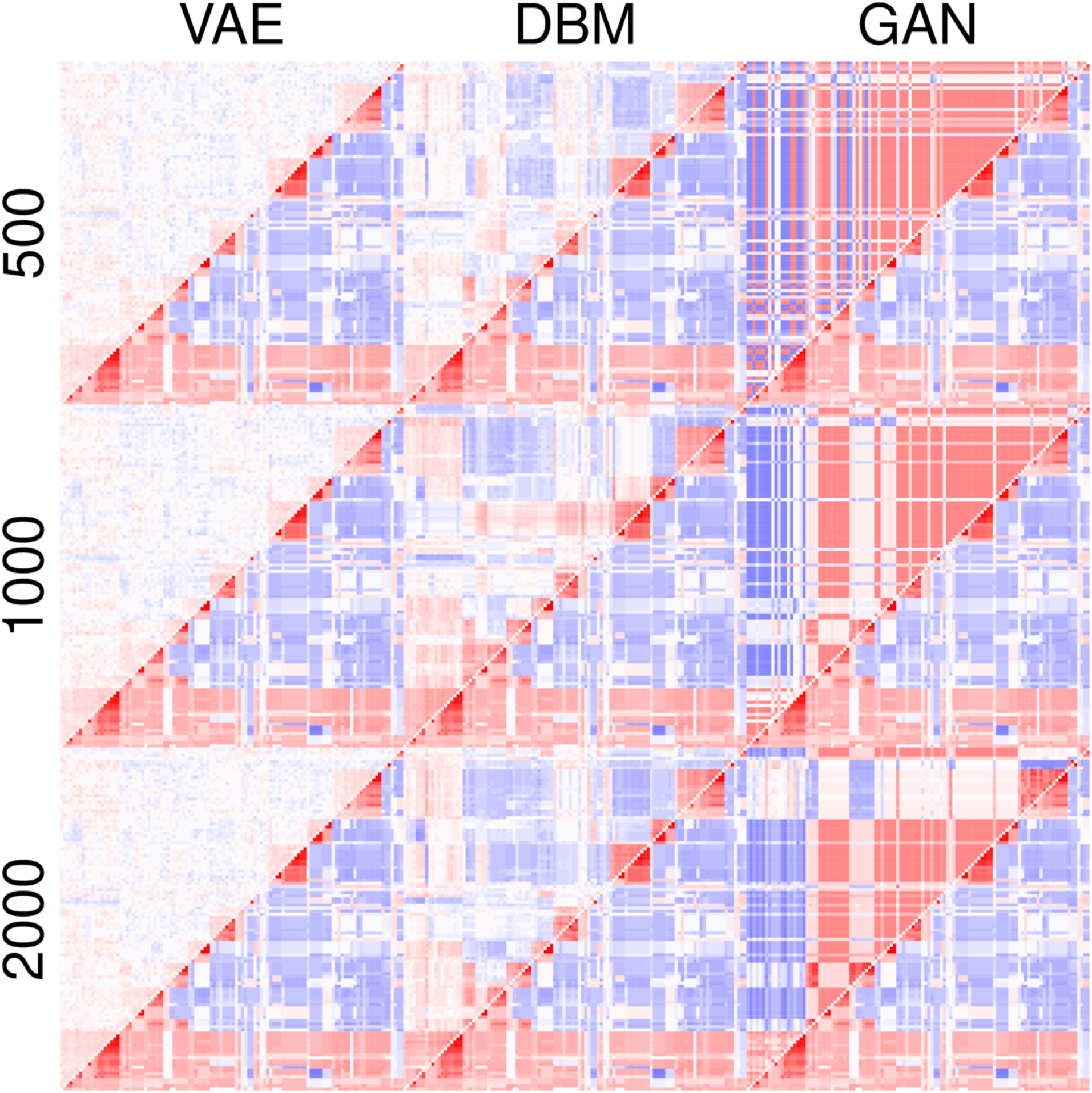
*Learned pair-wise log odds ratios in the HLA-B data. Figure shows pair-wise log odds ratios computed from 1000 synthetic observations, sampled from VAEs, DBMs and GANs trained with* 500, 1000 *and* 2000 *haplotypes. Log odds ratios computed from 36.8% of the 5008 real haplotypes, not used for training, are shown in the lower triangular of each log odds ratio matrix for sake of comparison. Red indicates positive log odds ratios while blue indicates negative log odds ratios*.

In contrast, the VAE and the DBM perform much better. However, as observed for the simulated non-conditional data, the VAE strongly under-estimates the log odds ratios and shows no large improvement with an increased amount of training data. The DBM benefits from increased training data size and overall delivers the best results on the level of MSE (Figure 4) and the visually graded log odds ratio pattern (Figure 5) as well as the generated synthetic haplotypes (Supplementary Figure 6).

### Conditional setting - Simulated data

We investigate how well GANs, specifically cGANs, and DBMs can infer the binary expression level of genes, conditional on the allelic variants of SNPs, where one allele affects a gene to be highly expressed and vice versa.

Similar to the non-conditional setting, we again train DBMs and cGANs with sub-samples of in total 10.000 observations and investigate the log odds ratios. As described in the methods section and shown on Supplementary Figure 1, we investigate the log odds ratios in an information carrying block, where there is a dependency between the SNP status and gene expression signal and in a non-information carrying block, lacking the dependency. In contrast to the non-conditional setting, the blocks do here relate to sub-sets of observations. In 50% of the observations, there is a dependency between the first 10 gene expression signals and the SNP status while this signal is absent in the remainder.

As in the non-conditional experiments, the cGAN has problems in picking up the signal in the information carrying block with lower amount of training data (Figure 6). However, with increased sample size, the results largely improve. The cGAN learns the distribution of log odds ratios in the non-information carrying block well (Supplementary Figure 7). Still, the synthetic observations, sampled from the DBM, do better reflect the correlation between SNP status and gene expression levels. This is also visible on the level of samples of the gene expression data conditional on the SNPs (Figure 7). While both architectures tend to capture the signal well from 1000 training observations onwards, cGAN tends to learn artifacts in the noise variables (see Figure 7).

**Figure 6:**
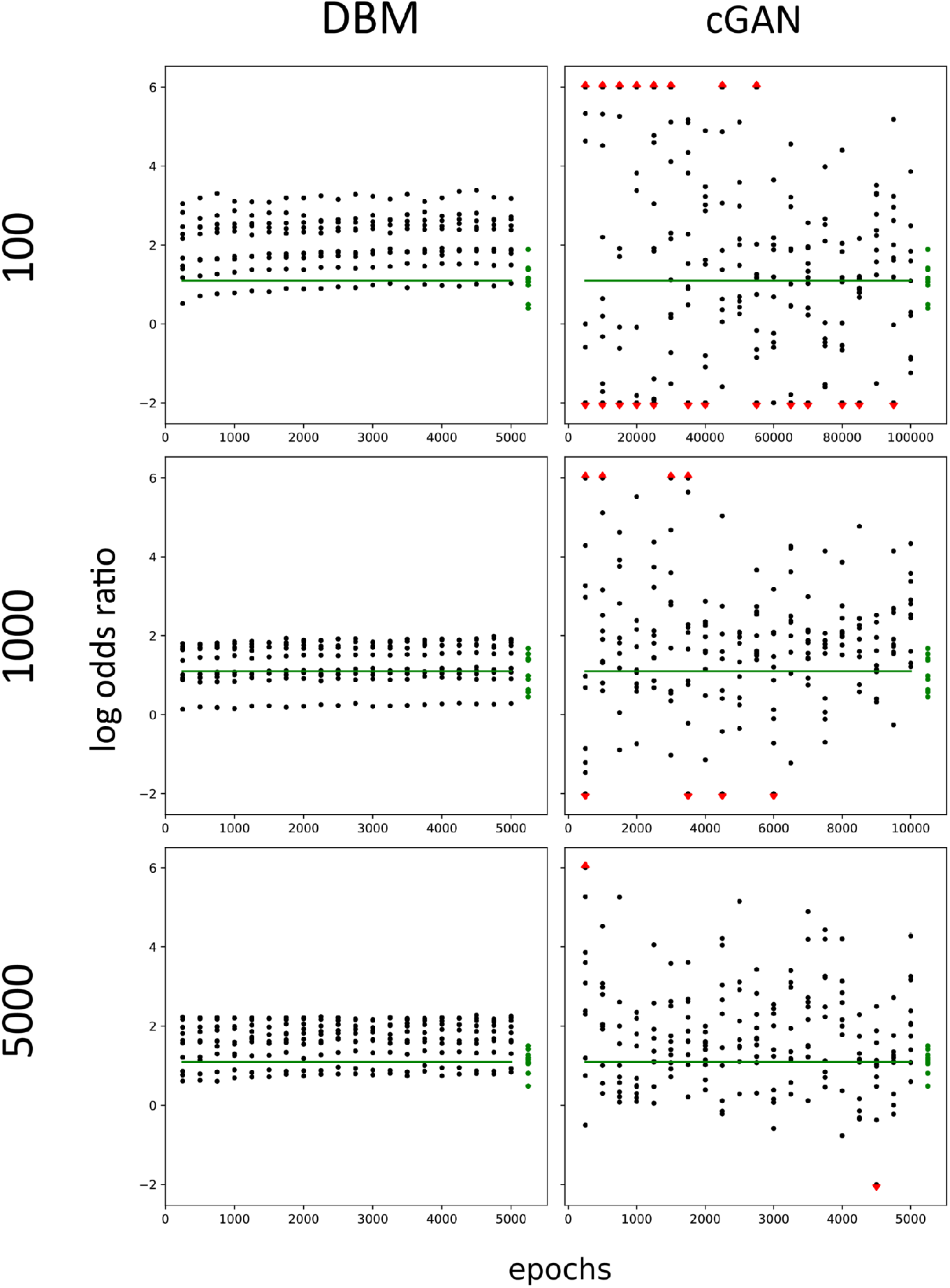
*Log odds ratios between genes and the states of SNPs in the **information carrying block**, computed during the training of DBMs and cGANs. DBMs and cGANs were trained on* 100, 1000 *and* 5000 *observations of simulated binary gene expression data, comprising the expression level of 50 genes. The same number of observations is provided for simulated SNP haplotype data, comprising 10 SNPs. Log odds ratios between SNPs and the affected genes, i.e. between SNP*_1_ *and Gene*_1_, *SNP*_2_ *and Gene*_2_ *and so on, are calculated* 20 *times during training. The green line shows the value of the theoretical truth according to the data generation procedure and the last entry of green dots shows the actual values of the training set*.

**Figure 7:**
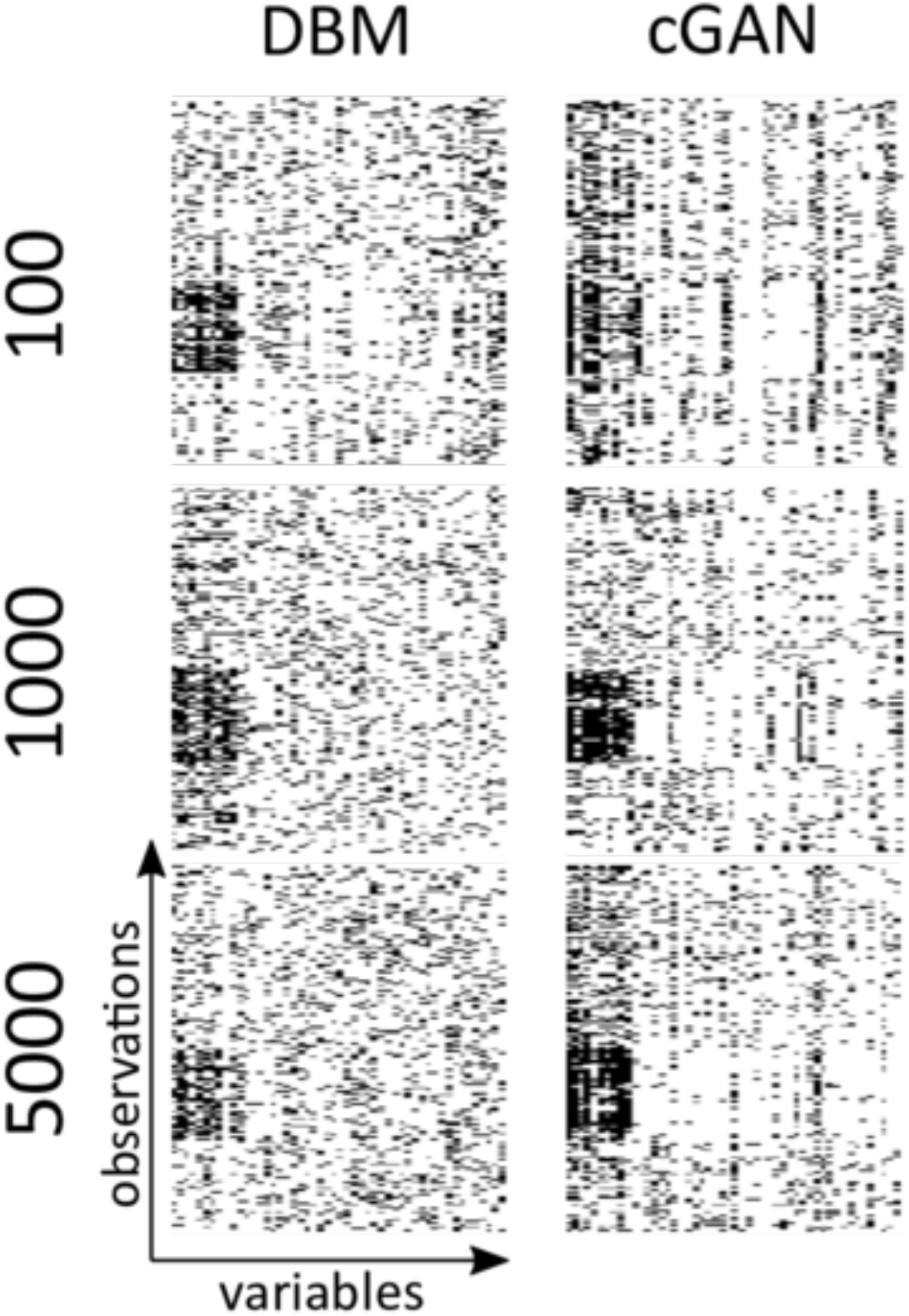
*Synthetic gene expression observations, sampled from trained DBMs and cGANs. DBMs and cGANs were trained on* 100, 1000 *and* 5000 *observations of simulated binary gene expression data, comprising the expression level of 50 genes. The same number of observations is provided for simulated SNP haplotype data, comprising 10 SNPs. Synthetic observations are generated by sampling the states of visual units representing the gene expression data, conditional on the states of the SNPs (DBM) or by propagating the activity of 10 randomly initialized noise variables together with the states of the 10 SNPs to the generator network and sampling from the units, representing the observed variables (cGAN). Synthetic observations are grouped based on their overall similarity by k-means clustering. Black indicates a 1 while white indicates a 0*.

In contrast to the non-conditional simulated scenario (Supplementary Figure 4), we observe no variability in the modeling performance for different data-sets in the conditional setting (Supplementary Figure 8).

## Discussion and Conclusions

We evaluated how well the joint distribution of binary data can be learnt by frequently employed deep generative modeling approaches, specifically VAEs, DBMs and GANs. We compared the approaches with a robust external measure, specifically the difference between the pair-wise log odds ratios computed from the empirical data, and the pair-wise log odds ratios computed from synthetic data, sampled from the models. We investigated the performance of the approaches with simulated data as well as real SNP data.

We observe, that DBMs generally learn the joint distribution of the simulated and real data well. While VAEs capture a signal even at a very small sample size, they fail in generating data that are visually similar to the training data. This is most likely due to their under-estimation of the strength of the signal in the data. Finally, given the investigated data, GANs are only capable to learn rather simple patterns at a rather large number of observations, used for training. We observe generally better results for the GAN in the conditional setting (cGAN). An explanation is, that the training objective is simplified by providing more information during training. This should result in a reduced parameter space which has to be explored. Another explanation is, that the signal was stronger in this scenario, compared with the non-conditional investigation.

There are two potential reasons, why DBMs perform better compared to GANs at rather small sample sizes and, compared to VAEs, generally better learn the magnitude of the signal. First, compared to DBMs, VAEs and GANs require to learn more parameters since they rely on feed-forward networks (see Figure 1). The second reason is related to the regularization which is applied during parameter optimization. In the VAE, we regularize the parameter updates by the Kullback-Leibler divergence between the Gaussian prior and the learnt latent posterior distribution. Here we have to a priori define the family of the distribution we assume for approximating the latent representation. Although being a reasonable choice in many applications, the employed normal distribution might not be appropriate for binary SNP data. In the DBM, on the other hand, we directly model the distribution of an observed binary variable, conditional on the distribution of other binary variables. Here the regularization is implicitly achieved by sampling from the joint distribution of observed and latent variables during training. In fact, it is known that MCMC techniques, as employed in the DBM, can generate asymptotically exact samples in contrast to variational techniques, as employed in the VAE [26]. In contrast, in the GAN, we do not have an intrinsic regularizer as in the DBM or the VAE. Instead regularizers and other hyper-parameters have to be carefully selected and this is not trivial, resulting in a performance, highly dependent on the employed method used for regularization [27].

The focus of this work is on the comparison of the three approaches in a small data setting close to real-life applications. We intended to simulate the scenario, in which a data scientist would try out the approaches using default settings, as it is usually the case with other applied biostatistics / bioinformatics methods. Consequently, all models were implemented “out of the box”. We intended to avoid intensive hyper-parameter tuning and employed rather simple neural network architectures. Thus results therefore might not show the full potential of the models. However, due to the poor performance of the GAN, we investigated several variants of GANs such as the Wasserstein GAN [28] but this did not result in considerably better performance.

With respect to our evaluation criterion, the pair-wise log odds ratio, one might argue, a limitation of our measure is, that more complex properties of the distributions are not captured. However we think that the pair-wise statistics do at least provide a lower bound for the quality of the learnt distribution, i.e. an approach that does not perform well here also won’t capture more complex structure. In fact, already this rather simple measure was generally in concordance with the subjective visual quality of samples. For instance the under-estimation of log odds ratios by the VAE (Figure 2) was also clearly observable in terms of an absence of visually clearly identifiable patterns in the synthetic data (Figure 3). Thus, the results provide also more general guidance of the use of deep generative approaches with categorical omics data.

## Funding

The work of MH has been supported by the Federal Ministry of Education and Research in Germany (BMBF) project “Generatives Deep-Learning zur explorativen Analyse von multimodalen Omics-Daten bei begrenzter Fallzahl” (GEMOLS: Generative deep-learning for exploratory analysis of muldimodal omics data with limited sample size, Fkz. 031L0250A). The work of SL has been supported by the BMBF in the MIRACUM project (Fkz. 01ZZ1801B).

## Notes

### Competing Interest Statement

The authors have declared no competing interest.

## Bibliography

1. Salakhutdinov R, Hinton G. Deep Boltzmann machines. Artificial intelligence and statistics 2009; 448–455

2. Kingma DP, Welling M. Auto-encoding variational bayes. arXiv preprint arXiv: 1312.6114 2013;

3. Goodfellow I, Pouget-Abadie J, Mirza M, et al. Generative adversarial nets. Advances in neural information processing systems 2014; 2672–2680

4. Beaulieu-Jones BK, Wu ZS, Williams C, et al. Privacy-preserving generative deep neural networks support clinical data sharing. Circulation: Cardiovascular Quality and Outcomes 2019; 12:e005122

5. Yelmen B, Decelle A, Ongaro L, et al. Creating artificial human genomes using generative models. bioRxiv 2019; 769091

6. Lopez R, Regier J, Cole MB, et al. Deep generative modeling for single-cell transcriptomics. Nature methods 2018; 15:1053

7. Rampášek L, Hidru D, Smirnov P, et al. Dr. VAE: Improving drug response prediction via modeling of drug perturbation effects. Bioinformatics 2019; 35:3743–3751

8. Hess M, Lenz S, Blätte TJ, et al. Partitioned learning of deep Boltzmann machines for SNP data. Bioinformatics 2017; 33:3173–3180

9. Eraslan G, Simon LM, Mircea M, et al. Single-cell RNA-Seq denoising using a deep count autoencoder. Nature communications 2019; 10:1–14

10. Eraslan G, Avsec Z, Gagneur J, et al. Deep learning: New computational modelling techniques for genomics. Nature Reviews Genetics 2019; 20:389–403

11. Choi E, Biswal S, Malin B, et al. Generating multi-label discrete patient records using generative adversarial networks. arXiv preprint arXiv:1703.06490 2017;

12. Sudmant PH, Rausch T, Gardner EJ, et al. An integrated map of structural variation in 2,504 human genomes. Nature 2015; 526:75

13. Theis L, Oord A van den, Bethge M. A note on the evaluation of generative models. arXiv preprint arXiv:1511.01844 2015;

14. Wang X, Ghasedi Dizaji K, Huang H. Conditional generative adversarial network for gene expression inference. Bioinformatics 2018; 34:i603–i611

15. Isola P, Zhu J-Y, Zhou T, et al. Image-to-image translation with conditional adversarial networks. Proceedings of the ieee conference on computer vision and pattern recognition 2017; 1125–1134

16. Mirza M, Osindero S. Conditional generative adversarial nets. arXiv preprint arXiv:1411.1784 2014;

17. Gauthier J. Conditional generative adversarial nets for convolutional face generation. Class Project for Stanford CS231N: Convolutional Neural Networks for Visual Recognition, Winter semester 2014; 2014:2

18. Ackley DH, Hinton GE, Sejnowski TJ. A learning algorithm for Boltzmann machines. Cognitive science 1985; 9:147–169

19. Salakhutdinov R, Mnih A, Hinton G. Restricted Boltzmann machines for collaborative filtering. Proceedings of the 24th international conference on machine learning 2007; 791–798

20. Abadi M, Barham P, Chen J, et al. TensorFlow: A system for large-scale machine learning. 12th {usenix} symposium on operating systems design and implementation ({osdi} 16) 2016; 265–283

21. Innes M. Flux: Elegant machine learning with Julia. Journal of Open Source Software 2018; 3:602

22. Bezanson J, Edelman A, Karpinski S, et al. Julia: A fresh approach to numerical computing. SIAM review 2017; 59:65–98

23. Lenz S, Hess M, Binder H. Unsupervised deep learning on biomedical data with BoltzmannMachines.jl. bioRxiv 2019; 578252

24. Snoke J, Raab GM, Nowok B, et al. General and specific utility measures for synthetic data. Journal of the Royal Statistical Society: Series A (Statistics in Society) 2018; 181:663–688

25. Bland JM, Altman DG. The odds ratio. BMJ 2000; 320:1468

26. Blei DM, Kucukelbir A, McAuliffe JD. Variational inference: A review for statisticians. Journal of the American statistical Association 2017; 112:859–877

27. Lucic M, Kurach K, Michalski M, et al. Are GANs created equal? A large-scale study. Advances in neural information processing systems 2018; 700–709

28. Arjovsky M, Chintala S, Bottou L. Wasserstein GAN. arXiv preprint arXiv:1701.07875 2017;

