## Supplementary Information for "Synthetic observations from deep generative models and binary omics data with limited sample size"

### Capability of deep generative architectures to model the distribution of categorical omics data under training data constraints - Supplementary Information

Jens Keienburg Frederic Boesel Stefan Lenz
Harald Binder Moritz Hess

June 5, 2020

2 **Log odds ratios learned by VAE, DBM and GAN from simulated SNP data - block 3.** VAEs, DBMs and GANs are trained on 100, 1000 and 5000 observations of simulated SNP haplotypes in 50 SNPs. Log odds ratios between negatively correlated SNPs (block 3, see figure 1) are calculated 20 times during training. Each dot represents the average of the log odds ratios between two SNPs computed from synthetic observations sampled from 200 generative models trained on different random sub-sets. The green line indicates the value of the theoretical truth according to the data generation procedure and the green dots indicate the actual values for the log odds ratios computed from the training set. Red arrows indicate values that are higher / lower than a log odds ratio of 1 / -2.5. . . . . . . . . . . . . . . 7

3 **Log odds ratios learned by VAE, DBM and GAN from simulated SNP data - block 4.** VAEs, DBMs and GANs are trained on 100, 1000 and 5000 observations of simulated SNP haplotypes in 50 SNPs. Log odds ratios between uncorrelated SNPs (block 4, see figure 1) are calculated 20 times during training. Each dot represents the average of the log odds ratios between two SNPs computed from synthetic observations sampled from 200 generative models trained on different random sub-sets. The green line indicates the value of the theoretical truth according to the data generation procedure and the green dots indicate the actual values for the log odds ratios computed from the training set. Red arrows indicate values that are higher / lower than a log odds ratio of 3 / -3. . . . . . . . . . . . . . . . . . . . . . . . . . 8

4 **Variability in training dependent on data set.** VAEs, DBMs and GANs are trained on 5000 observations of simulated SNP haplotypes in 50 SNPs. Log odds ratios between positively correlated SNPs (block 1, see figure 1) are calculated 20 times during training. Each dot represents the average of the log odds ratios between two SNPs computed from synthetic observations sampled from 200 generative models trained on different random sub-sets. Random sub-sets are drawn from four different data-sets (dataset 1 to 4). The green line indicates the value of the theoretical truth according to the data generation procedure and the green dots indicate the actual values for the log odds ratios computed from the training set. . . . . . . . . . . . . . . . . . . . 9
5 **Investigated haplotypes from the HLA-B locus.** The upper panel shows log odds ratios of 110 SNPs and the lower panel shows the 5008 haplotypes, used to compute the log odds ratios. Red indicates positive log odds ratios while blue indicates negative log odds ratios. . . . . . . . . . . . . . . . . . . . . . . . . . . . . . 10
6 **Synthetic data, sampled from deep generative models.** Models were trained on 63.2% of the 5008 haplotypes in the 1000 genomes data. 110 SNPs in the HLA-B locus are investigated. 1000 synthetic observations are shown for each sample size and each model. . . . . . . . . . . . . . . . . . . . . . . . . . . . . . 11
7 **Log odds ratios between genes and the states of SNPs in the non-information carrying block, computed during the training of DBMs and cGANs.** DBMs and cGANs were trained on 100, 1000 and 5000 observations of simulated binary gene expression data, comprising the expression level of 50 genes. The same number of observations is provided for simulated SNP haplotype data, comprising 10 SNPs. Log odds ratios between SNPs and the corresponding, associated gene expression variables are calculated 20 times during training. The information carrying data holds positively correlated features. The green line shows the value of the theoretical truth according to the data generation procedure and the last entry of green dots shows the actual values of the training set. . . . . . . . . . . . . . . . . . . . . . . . . 12
8 **Variability in training dependent on data set - information carrying block.** DBMs and cGANs are trained on 5000 observations of simulated binary gene expression data, comprising the expression level of 50 genes. The same number of observations is provided for simulated SNP haplotype data, comprising 10 SNPs. Log odds ratios between SNPs and gene expression are calculated 20 times during training. Random sub-sets are drawn from four different data-sets (dataset 1 to 4). The green line shows the value of the theoretical truth according to the data generation procedure and the last entry of green dots shows the actual values of the training set. . . . . . . . . . . . . . . . . . . . . . . . . 13

### Architectures

Here we provide details about the employed architectures. All abbreviations relate to figure 1 in the main manuscript.

#### Simulated non-conditional

##### VAE

- learning-rate = 0.00001
- batch-size = 32

| Layer | Nodes | Activation function |
|---|---|---|
| h_enc | 30 | Tangens hyperbolicus |
| mu | 10 | identity |
| sigma | 10 | identity |
| h_dec | 30 | Tangens hyperbolicus |
| xhat | 50 | sigmoid |

##### DBM

For sample sizes of 100 to 5000, the layer-wise pre-training is conducted for 4000 to 80 epochs with a learning rate of 0.001. Training of the overall DBM made use of the mean-field-approximation technique, where the learning rate was set to 0.05 for all DBM experiments.

| Layer | Nodes | Activation function |
|---|---|---|
| x | 50 | sigmoid |
| h_1 | 50 | sigmoid |
| h_2 | 10 | sigmoid |

##### GAN

- learning-rate = 0.00001
- batch-size = 32

| Layer | Nodes | Activation function |
|---|---|---|
| z | 10 | |
| h_gen | 30 | Leaky Relu |
| xhat | 50 | sigmoid |
| x , xhat | 50 | |
| h_disc | 25 | Leaky Relu |
| d(x , xhat) | 1 | sigmoid |

#### Empirical SNP data

##### VAE

- learning-rate = 0.001
- batch-size = 10

| Layer | Nodes | Activation function |
|---|---|---|
| h_enc | 30 | Tangens hyperbolicus |
| mu | 10 | identity |
| sigma | 10 | identity |
| h_dec | 30 | Tangens hyperbolicus |
| xhat | 50 | sigmoid |

##### DBM

We employed a learningrate for the layer-wise pre-training and fine-tuning of 0.0001 and performed fine-tuning for 10 epochs. We tuned the number of epochs of pre-training. In total, we monitored the training process over 8000 epochs.

- batch-size = 10

| Layer | Nodes | Activation function |
|---|---|---|
| x | 50 | sigmoid |
| h_1 | 50 | sigmoid |
| h_2 | 10 | sigmoid |

**GAN**

- learning-rate = 0.001
- batch-size = 10

| Layer | Nodes | Activation function |
|---|---|---|
| z | 10 | |
| h_gen | 30 | Leaky Relu |
| xhat | 50 | sigmoid |
| x , xhat | 50 | |
| h_disc | 25 | Leaky Relu |
| d(x , xhat) | 1 | sigmoid |

#### Simulated conditional

##### DBM

The layer-wise pre-training with learning rate 0.002 has been applied for 4000 to 80 epochs for the sample sizes 100 to 5000. The overall DBM was trained with a learning rate of 0.003 for 5000 epochs.

| Layer | Nodes | Activation function |
|---|---|---|
| x | 60 | sigmoid |
| h_1 | 30 | sigmoid |
| h_2 | 10 | sigmoid |
| h_3 | 2 | sigmoid |

##### cGAN

- learning-rate = 0.0001
- batch-size = 32

| Layer | Nodes | Activation function |
|---|---|---|
| z | 10 | |
| h_gen | 20 | Leaky Relu |
| xhat | 50 | sigmoid |
| x , xhat | 50 | |
| h_disc1 | 60 | Leaky Relu |
| h_disc2 | 40 | Leaky Relu |
| h_disc3 | 20 | Leaky Relu |
| d(x , xhat \|c) | 1 | sigmoid |

#### Supplementary figures

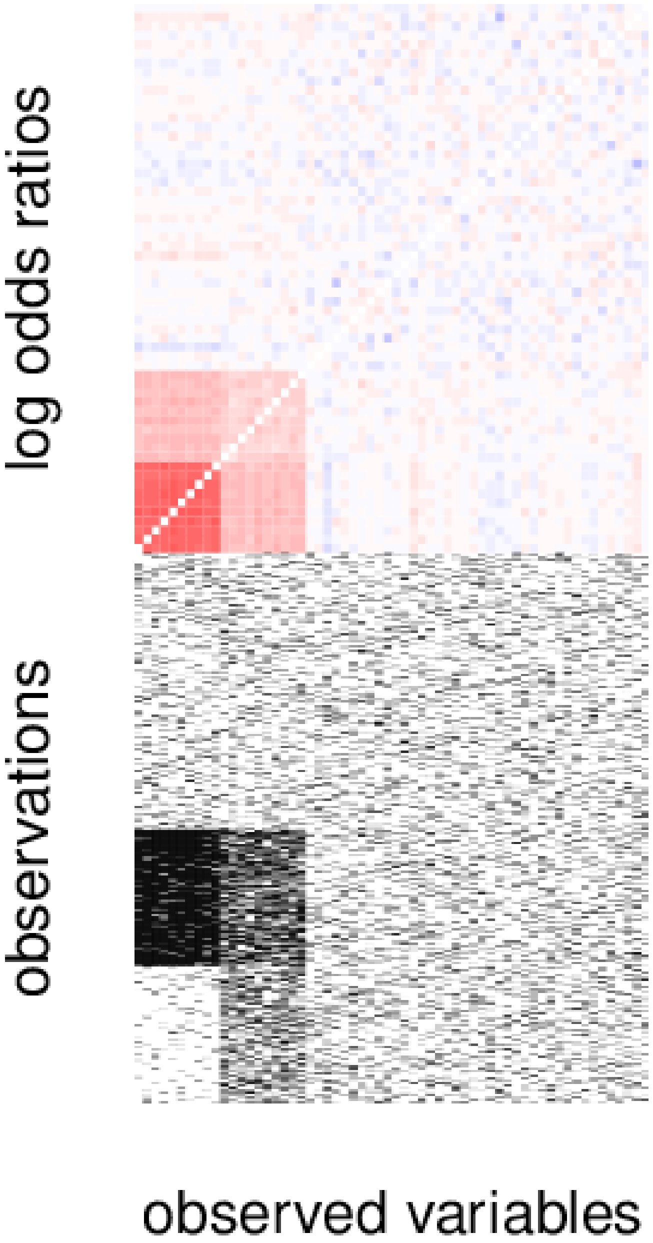

Supplementary Figure 1: **Simulated SNP and gene expression data for evaluating DBMs and cGANs in the conditional setup.** The first 10 observed variables are the SNPs and the second 10 variables represent the gene expression of genes that are affected by the SNP status. The upper 50% of observations represent the non-information carrying block while the lower 50% represent the information carrying block.

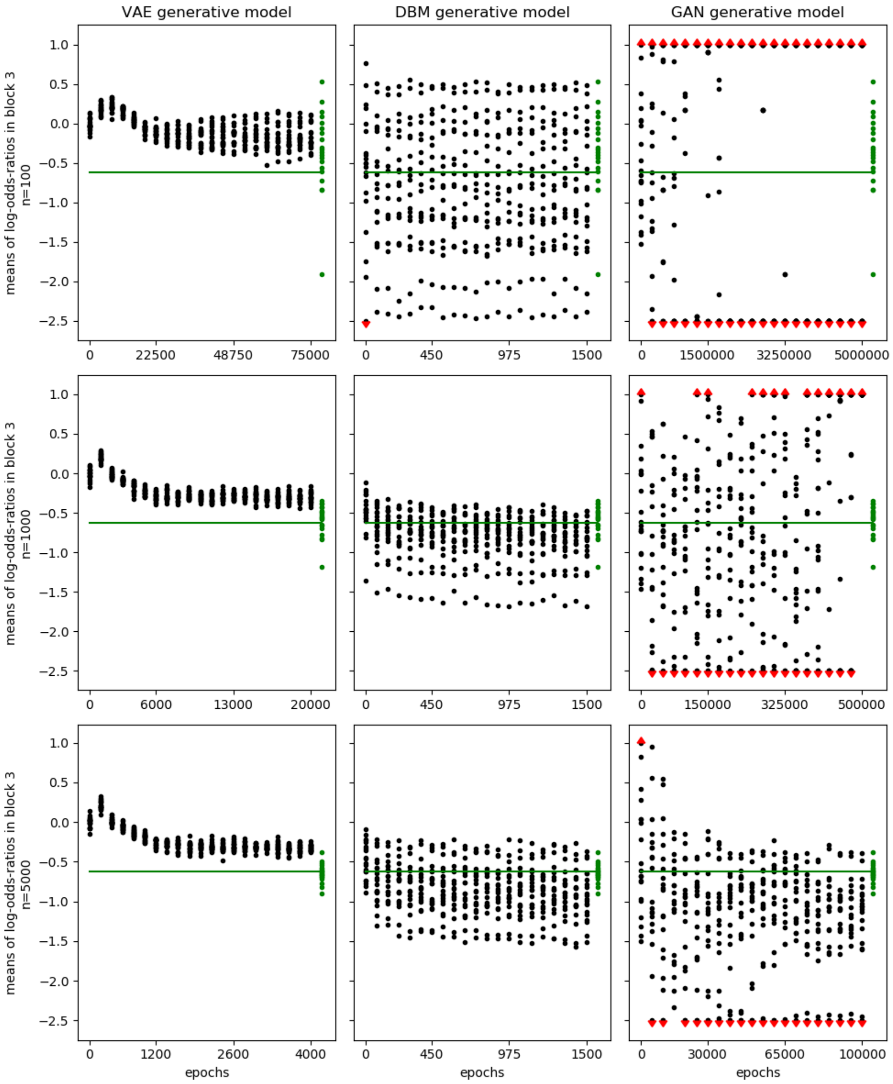

Supplementary Figure 2: **Log odds ratios learned by VAE, DBM and GAN from simulated SNP data - block 3.** VAEs, DBMs and GANs are trained on 100, 1000 and 5000 observations of simulated SNP haplotypes in 50 SNPs. Log odds ratios between negatively correlated SNPs (block 3, see figure 1) are calculated 20 times during training. Each dot represents the average of the log odds ratios between two SNPs computed from synthetic observations sampled from 200 generative models trained on different random sub-sets. The green line indicates the value of the theoretical truth according to the data generation procedure and the green dots indicate the actual values for the log odds ratios computed from the training set. Red arrows indicate values that are higher / lower than a log odds ratio of 1 / -2.5.

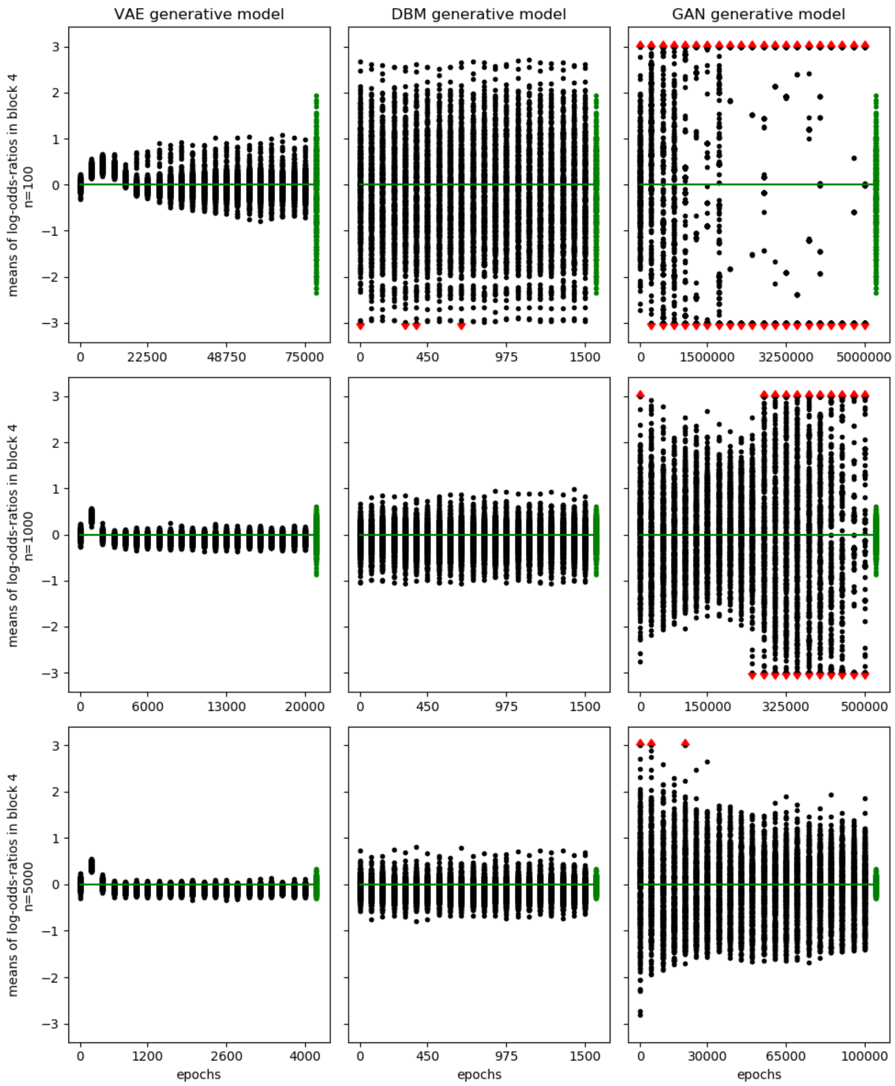

Supplementary Figure 3: **Log odds ratios learned by VAE, DBM and GAN from simulated SNP data - block 4.** VAEs, DBMs and GANs are trained on 100, 1000 and 5000 observations of simulated SNP haplotypes in 50 SNPs. Log odds ratios between uncorrelated SNPs (block 4, see figure 1) are calculated 20 times during training. Each dot represents the average of the log odds ratios between two SNPs computed from synthetic observations sampled from 200 generative models trained on different random sub-sets. The green line indicates the value of the theoretical truth according to the data generation procedure and the green dots indicate the actual values for the log odds ratios computed from the training set. Red arrows indicate values that are higher / lower than a log odds ratio of 3 / -3.

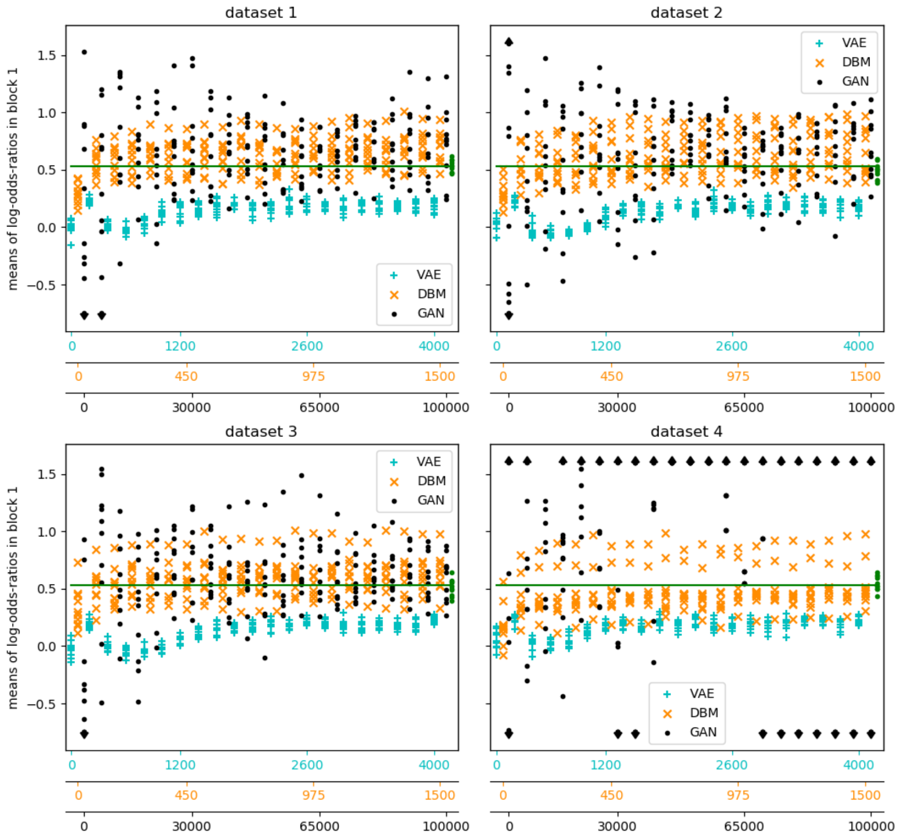

Supplementary Figure 4: **Variability in training dependent on data set.** VAEs, DBMs and GANs are trained on 5000 observations of simulated SNP haplotypes in 50 SNPs. Log odds ratios between positively correlated SNPs (block 1, see figure 1) are calculated 20 times during training. Each dot represents the average of the log odds ratios between two SNPs computed from synthetic observations sampled from 200 generative models trained on different random sub-sets. Random sub-sets are drawn from four different data-sets (dataset 1 to 4). The green line indicates the value of the theoretical truth according to the data generation procedure and the green dots indicate the actual values for the log odds ratios computed from the training set.

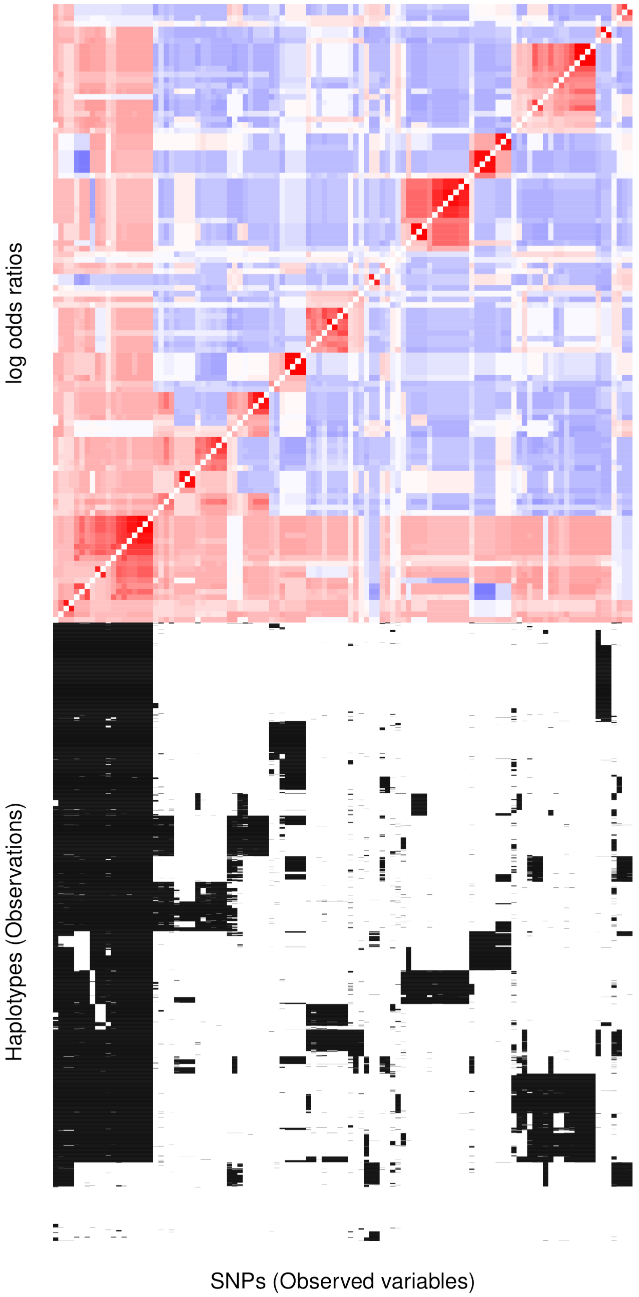

Supplementary Figure 5: **Investigated haplotypes from the HLA-B locus.** The upper panel shows log odds ratios of 110 SNPs and the lower panel shows the 5008 haplotypes, used to compute the log odds ratios. Red indicates positive log odds ratios while blue indicates negative log odds ratios.

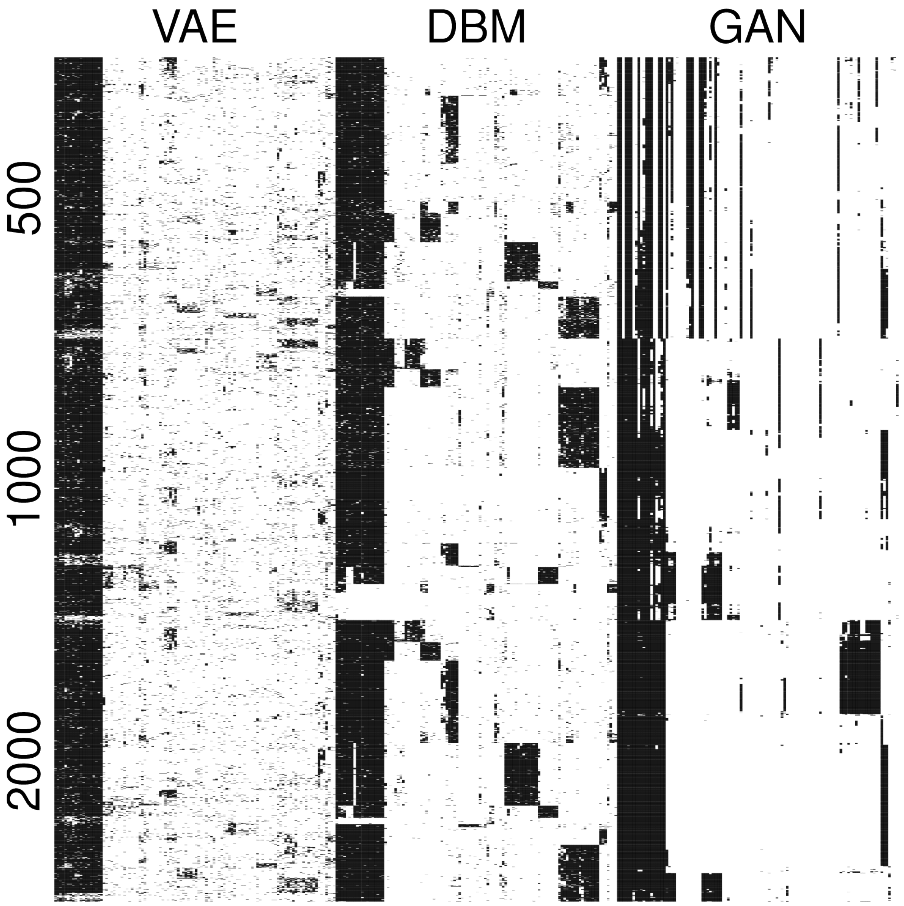

Supplementary Figure 6: **Synthetic data, sampled from deep generative models.** Models were trained on 63.2% of the 5008 haplotypes in the 1000 genomes data. 110 SNPs in the HLA-B locus are investigated. 1000 synthetic observations are shown for each sample size and each model.

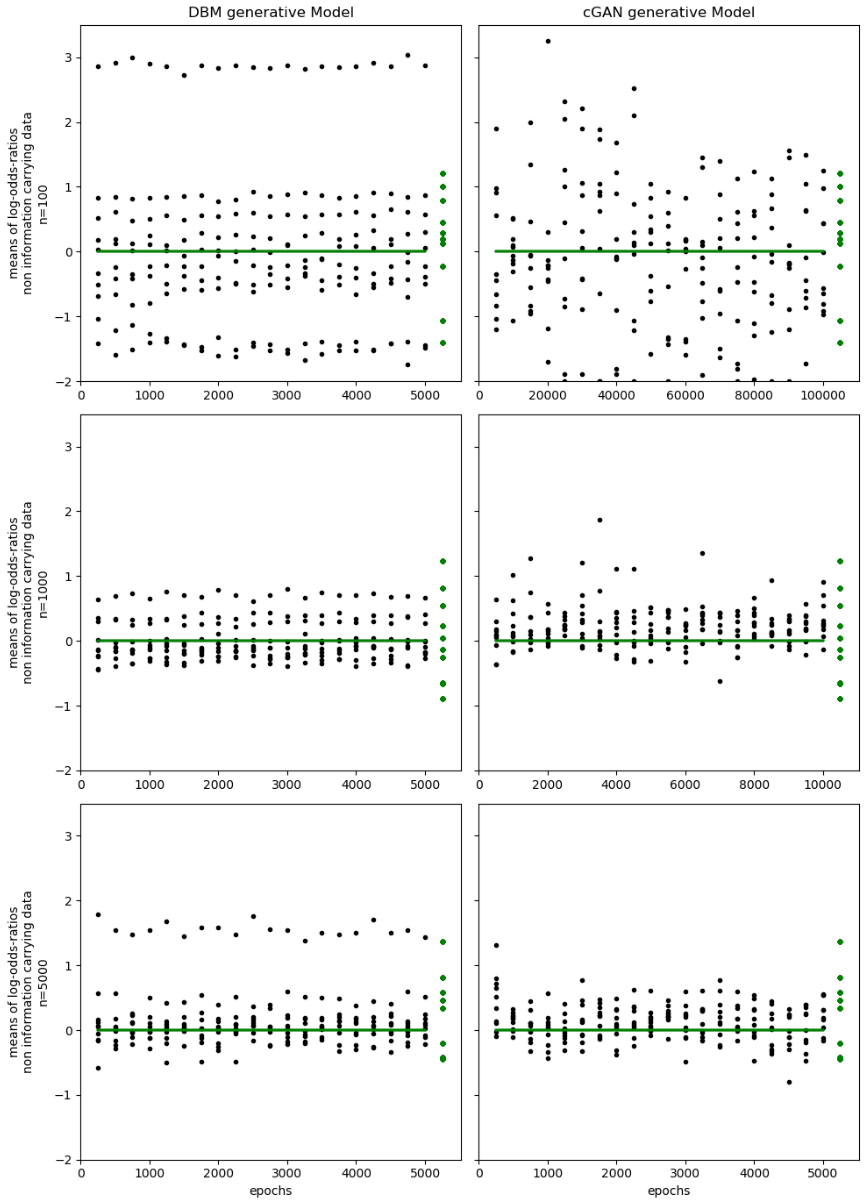

Supplementary Figure 7: **Log odds ratios between genes and the states of SNPs in the non-information carrying block, computed during the training of DBMs and cGANs.** DBMs and cGANs were trained on 100, 1000 and 5000 observations of simulated binary gene expression data, comprising the expression level of 50 genes. The same number of observations is provided for simulated SNP haplotype data, comprising 10 SNPs. Log odds ratios between SNPs and the corresponding, associated gene expression variables are calculated 20 times during training. The information carrying data holds positively correlated features. The green line shows the value of the theoretical truth according to the data generation procedure and the last entry of green dots shows the actual values of the training set.

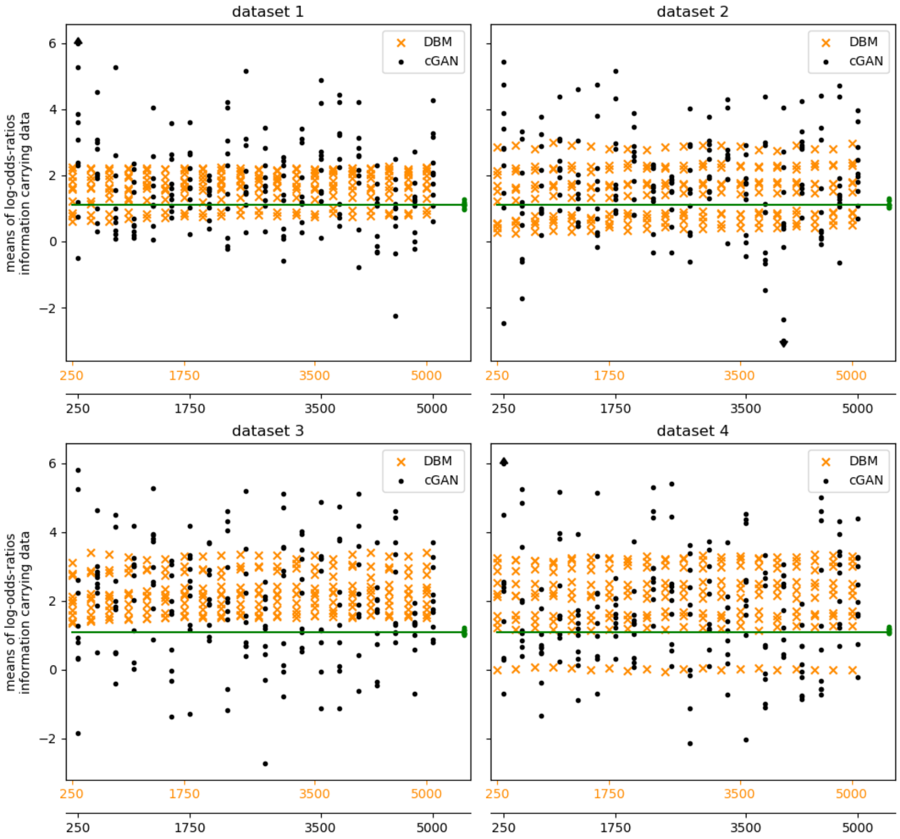

Supplementary Figure 8: **Variability in training dependent on data set - information carrying block.** DBMs and cGANs are trained on 5000 observations of simulated binary gene expression data, comprising the expression level of 50 genes. The same number of observations is provided for simulated SNP haplotype data, comprising 10 SNPs. Log odds ratios between SNPs and gene expression are calculated 20 times during training. Random sub-sets are drawn from four different data-sets (dataset 1 to 4). The green line shows the value of the theoretical truth according to the data generation procedure and the last entry of green dots shows the actual values of the training set.
